# Social learning of a challenging two-step action sequence fulfils key criteria of cumulative culture in an insect

**DOI:** 10.1101/2023.08.29.555358

**Authors:** Alice D. Bridges, Amanda Royka, Tara Wilson, Lars Chittka

## Abstract

Culture in non-human animals refers to behaviour that is socially learned and persists within a population over time. Human culture is notable in that it is *cumulative*: new innovations have built on previous ones over thousands of years. As a result, what is acquired via social learning often goes far beyond the capacity of any individual to independently discover it during their lifetime^1–3^. To date, no previous study has convincingly demonstrated this phenomenon in a non-human animal. Here, we show that bumblebees can learn from a trained demonstrator to open a novel, 3D-printed two-step puzzle box to obtain food rewards, even though they fail to do so independently. Experimenters were unable to train demonstrators to perform the unrewarded first step of the behaviour without providing a temporary reward linked to this action: the reward then being removed during later stages of training. However, a third of naïve observers learned to open the two-step box from these demonstrators, without ever having been rewarded after the first step. This suggested that social learning might permit the acquisition of behaviours too complex to “re-innovate” via individual learning. Furthermore, naïve bees failed to open the box despite extended exposure over the course of 12 days. The temporal and spatial distance of the unrewarded first step from the reward appeared to inhibit acquisition of two-step box-opening via individual associative learning, but this limitation was overcome by the presence of a trained conspecific. To our knowledge, these results make bumblebees the first animal to demonstrate the ability to socially learn a behaviour that is beyond their ability to innovate individually. This finding challenges the prevailing opinion of the field, which generally considers cumulative culture, which is built on this capacity, to be unique to humans.

## Introduction

Culture in animals can be broadly conceptualised as the sum of a population’s behavioural traditions, which in turn are defined as behaviours that are transmitted via social learning and that persist within a population over time^4^. While culture was once thought to be exclusive to humans and a key explanation of our own evolutionary success, the existence of non-human cultures that change over time is no longer controversial. Changes in the songs of Savannah sparrows^5^ and humpback whales^6–8^ have been documented over decades. The sweet-potato washing behaviour of Japanese macaques has also undergone several distinctive modifications since its inception at the hands of “Imo”, a juvenile female, in 1953^9^. Imo’s initial behaviour involved dipping a potato in a freshwater stream and wiping sand off with her spare hand, but within a decade had evolved to include repeatedly washing in sea water in-between bites rather than in freshwater; potentially to enhance their flavour. By the 1980s, a range of variations had appeared among macaques, including stealing already-washed potatoes from conspecifics, and digging new pools in secluded areas to wash potatoes without being seen by scroungers^9–11^. Likewise, the “wide”, “narrow” and “stepped” designs of pandanus tools, which are fashioned from torn leaves by New Caledonian crows and used to fish grubs from logs, appear to have diverged from a single point of origin^12^. In this manner, cultural evolution can result in both the accumulation of novel traditions, and the accumulation of modifications to these traditions in turn. However, the limitations of non-human cultural evolution remain the subject of debate. Most notably, it remains unclear whether modifications can accumulate to the point where the final form of a behaviour is too complex for any individual to innovate itself, yet can still be acquired by that same individual via social learning from a knowledgeable conspecific. This threshold, beyond which culture can be considered substantially “cumulative”, is widely considered to represent a fundamental distinction between humans and non-humans.

It is clearly true that humans are a uniquely encultured species. Almost everything we do relies on knowledge or technology that has taken many generations to build. No one human being could possibly manage, within their own lifetime, to split the atom by themselves from scratch. They could not even conceive of doing so without centuries of accumulated scientific knowledge. The existence of this “cumulative culture” relies on the “ratchet” concept; whereby traditions are retained within a population with sufficient fidelity to allow improvements to accumulate^1–3^. This has been argued to require so-called “higher order” forms of social learning, such as true imitation^13^ or teaching^14^, and these in turn have been argued to be exclusive to humans. But if we strip the definition of cumulative culture back to its bare bones, for a behaviour to be considered *cumulative*, it must fulfil three requirements. It must build on an already-established behaviour among individuals of a population, naïve members of the population must be able to acquire the complete behaviour from knowledgeable individuals via social learning, and they must be unable to arrive at the same behaviour solely through innovation.

Bumblebees (*Bombus terrestris*) are social insects known to be capable of acquiring complex, non-natural behaviours via social learning in a laboratory setting, such as string-pulling^15^ and ball-rolling to gain rewards^16^. In the latter case, they were even able to improve on the behaviour of their original demonstrator. More recently, when challenged with a two-option puzzle-box task and a paradigm allowing learning to diffuse across a population (a gold-standard of cultural transmission experiments^17^), they were found to acquire *and* maintain arbitrary variants of this behaviour from trained demonstrators^18^. However, these previous investigations involved the acquisition of a behaviour that each bee could also have innovated independently. Indeed, some naïve individuals were able to open the puzzle box, pull strings and roll balls without demonstrators^15,16,18^. Thus, to determine whether bumblebees could acquire a behaviour through social learning that they could not innovate independently, we developed a novel two-step puzzle box (Fig. 1A). This design was informed by a “lockbox” task developed to assess problem solving in Goffin’s cockatoos^19^. Here, cockatoos were challenged to open a box that was sealed with five inter-connected “locks” that had to be opened sequentially, with no reward for opening any but the final lock. Our hypothesis was that this degree of temporal and spatial separation between performing the first step of the behaviour and the reward would make it very difficult, if not impossible, for a naïve bumblebee to form a lasting association between this necessary initial action and the final reward. Even if a bee opened the two-step box independently through repeated, non-directed probing, as observed with our previous box^18^, if no association formed between the combination of the two pushing behaviours and the reward, this behaviour would be unlikely to be incorporated into an individual’s repertoire. If, however, a bee was able to learn this multi-step box-opening behaviour when exposed to a skilled demonstrator, this would suggest that bumblebees can acquire behaviours socially that lie beyond their capacity for individual innovation.

**Figure 1.**
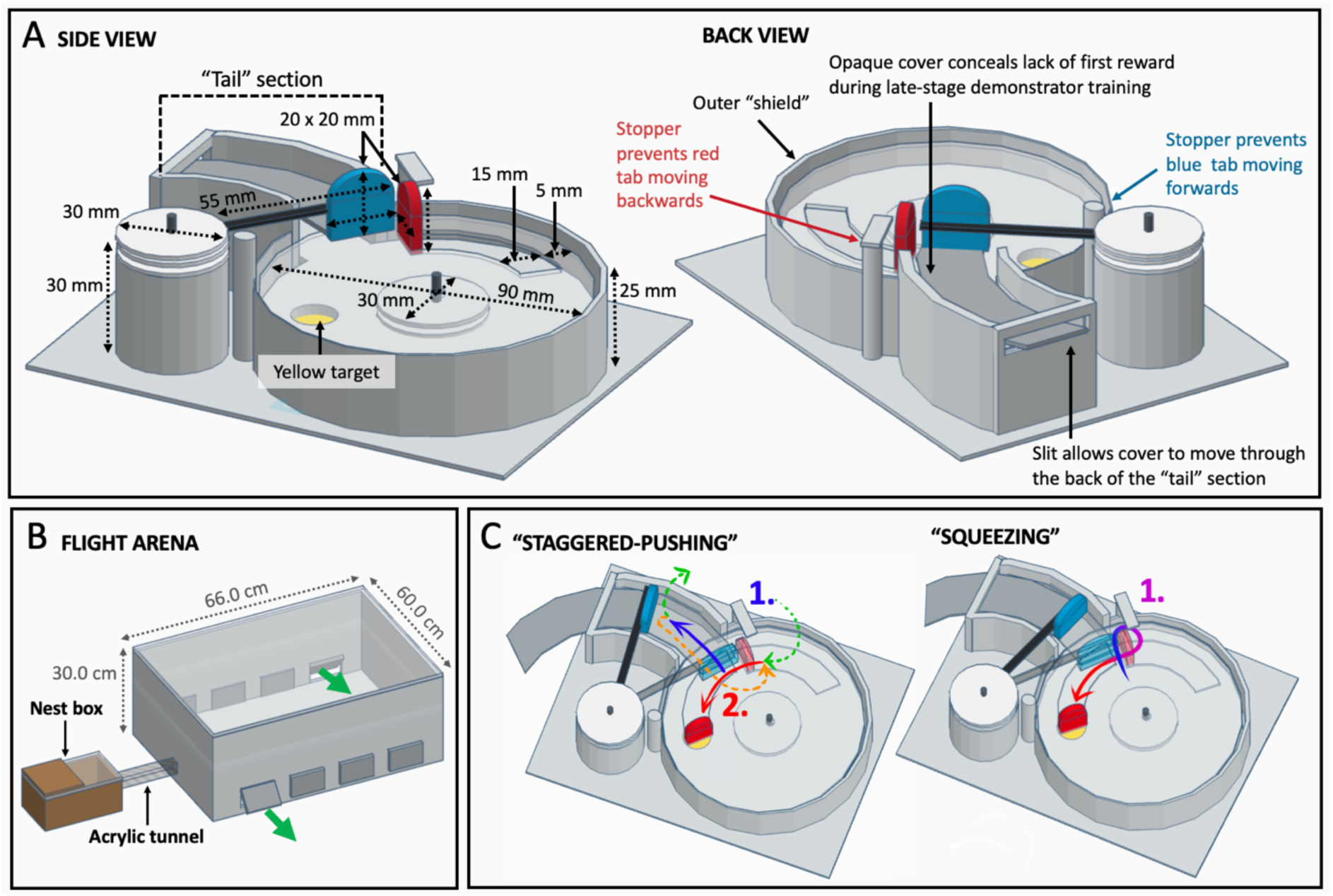
Two-step puzzle box design and experimental set-up. **(A) Puzzle box design.** Box bases were 3D-printed to ensure consistency. The reward (50% w/w sucrose solution, placed on a yellow target) was inaccessible unless the red tab was pushed, rotating the lid anti-clockwise around a central axis, and the red tab could not move unless the blue tab was first pushed out of its path. See Supplementary Information for a full description of the box design elements. **(B) Experimental set-up.** The flight arena was connected to the nest box via an acrylic tunnel, and flaps cut into the side allowed the removal and replacement of puzzle boxes during the experiment. The sides were lined with bristles to prevent bees escaping. **(C) Alternative action patterns for opening the box.** The “staggered-pushing” technique is characterised by two distinct pushes (1., blue arrow and 2., red arrow), divided by either flying (green arrows) or walking in a loop around the inner side of the red tab (orange arrow). The “squeezing” technique is characterised by an oftentimes-single, unbroken movement starting at the point where the blue and red tabs meet and pushing through; squeezing between the outer side of the red tab and the outer shield, and making a tight turn to immediately push against the red tab

## Results

### Naïve observers acquired two-step box-opening from trained demonstrators in a paired dyad experimental setting

The two-step puzzle box (Fig. 1A) relied on the same principles as our previous single-step, two-option puzzle box^18^. To access a 50% w/w sucrose solution reward, placed on a yellow target, a clear lid had to be rotated around a central axis by pushing against a coloured tab: in this case, by pushing a red tab anticlockwise. However, this tab could only move forward if another blue tab was pushed out of its path. A sample video of a trained demonstrator opening the two-step box can be found here.

The paired dyad experiments were conducted in a specially constructed flight arena, attached to a colony’s nest box, where all bees not currently undergoing training or testing were confined (Fig. 1B). In these experiments, a pair of bees, including one trained demonstrator and one naïve observer, were allowed to forage together on three closed puzzle boxes (each filled with 20 μl 50% w/w sucrose solution) for 30-40 sessions, with unrewarded learning tests given to the observer in isolation after 30, 35 and 40 joint sessions. If an observer passed a learning test, it immediately proceeded to 10 solo foraging sessions in the absence of the demonstrator. The 15 demonstrator and observer combinations used for the paired dyad experiments are listed in Table 1, with some demonstrators being used for multiple observers. Of the 15 observers, five passed the unrewarded learning test, with three of these doing so on the first attempt and the remaining two on the third. This relatively low number reflected the difficulty of the task, but the fact that any observers acquired two-step box-opening at all confirmed that this behaviour could be socially learned.

**Table 1.** Demonstrator and observer combinations, with outcomes.

| Demonstrator ID | Demonstrator technique | Demo opening incidence | Demo opening index | Observer ID | Outcome | Test passed | Solo foraging phase box-opening incidence | Of which were "squeezing" |
| --- | --- | --- | --- | --- | --- | --- | --- | --- |
| B-8 | Squeezing | 132 | 4.4 | A-12 | ✓ | 1 | n/a<sup>#</sup> | n/a<sup>#</sup> |
| B-8 | Squeezing | 137 | 3.4 | A-17 | ✓ | 3 | n/a<sup>#</sup> | n/a<sup>#</sup> |
| A-17* | Squeezing | 215 | 7.2 | A-24 | ✓ | 1 | 9 | 9 |
| A-17* | Squeezing | 187 | 6.2 | A-6 | ✓ | 1 | 13 | 9 |
| A-24* | Squeezing | 200 | 5.0 | A-37 | ✗ | n/a | n/a | n/a |
| A-24* | Squeezing | 222 | 5.6 | A-39 | ✗ | n/a | n/a | n/a |
| A-24* | Squeezing | 181 | 4.5 | C-42 | ✓ | 3 | 1 | 1 |
| C-15 | Staggered-pushing | 158 | 3.9 | C-26 | ✗ | n/a | n/a | n/a |
| C-26* | Squeezing | 72 | 1.8 | C-19 | ✗ | n/a | n/a | n/a |
| D-3 | Staggered-pushing | 227 | 5.7 | D-11 | ✗ | n/a | n/a | n/a |
| D-1 | Squeezing | 206 | 5.2 | D-77 | ✗ | n/a | n/a | n/a |
| D-72 | Squeezing | 103 | 3.4 | D-76 | ✗ | n/a | n/a | n/a |
| D-23 | Staggered-pushing | 258 | 6.8 | D-32 | ✗ | n/a | n/a | n/a |
| D-23 | Staggered-pushing | 315 | 7.9 | D-42 | ✗ | n/a | n/a | n/a |
| D-23 | Staggered-pushing | 234 | 7.8 | D-48 | ✗ | n/a | n/a | n/a |

\*Individual was originally used as an observer. In an effort to reduce training time (~2 days per entirely naive bee), these bees then went through stepwise demonstrator training and were used as demonstrators for other observers when they passed the unrewarded demonstrator learning test. The letters in demonstrator and observer IDs refer to the colony that bee originated from (A, B, C, or D), and the numbers refer to the individual tag attached to the thorax. <sup>#</sup>Bee was used to trial solo foraging bout protocol; therefore results are excluded due to being incomparable.

The post-learning test solo foraging sessions were designed to further confirm observer acquisition of two-step box-opening. Each session lasted up to 10 min, but 50 μl 50% sucrose solution was placed on the yellow target in each box: as *Bombus terrestris* foragers have been found to collect 60-150 μl sucrose solution per foraging trip depending on their size, this meant that each bee could reasonably be expected to open two boxes per session^20^. While all bees who proceeded to the solo foraging stage repeated two-step box opening, confirming their status as learners, only two individuals (A-24 and A-6; Table 1) met the criterion to be classified as *proficient* learners (i.e. they opened ≥10 boxes). This was the same threshold applied to learners in our previous work with the single-step two-option box^18^. However, it should be noted that learners from our present study had comparatively limited post-learning exposure to the boxes (a total of 100 min on one day) compared with our previous work. Proficient learners from our single-step puzzle-box experiments typically attained proficiency over several days of foraging, and had access to boxes for 180 min each day for 6-12 days^18^. Thus, these comparatively low numbers are perhaps unsurprising.

### Alternative action-patterns arose among two-step box-opening demonstrators, and impacted observer acquisition of the behaviour

Two different methods of opening the two-step puzzle box were observed among the trained demonstrators during the paired dyad experiments, and were termed “staggered-pushing” and “squeezing” (Fig. 1C). Of these techniques, “squeezing” typically resulted in the blue tab being pushed less far compared with “staggered-pushing”, often only just enough to free the red tab, and the red tab often shifted forward as the bee squeezed between this and the outer shield. Among demonstrators, the “squeezing” technique was more common, being adopted as the primary technique by 6/9 individuals (Table 1). Thus, 10/15 observers were paired with a “squeezing” demonstrator.

While not all observers paired with “squeezing” demonstrators learned to open the two-step box (5/10 succeeded), all observers paired with “staggered-pushing” demonstrators (n=5) failed to learn two-step box-opening. This discrepancy was not due to the number of demonstrations being received by the observers: there was no difference in the number of boxes opened by “squeezing” demonstrators compared with “staggered-pushing” demonstrators when the number of joint sessions was accounted for (unpaired t-test, t=-2.015, p=0.065; Table 2). This may have been because the “squeezing” demonstrators often performed their “squeezing” action several times, looping around the red tab, which lengthened the total duration of the behaviour despite the blue tab being pushed less than during “staggered-pushing”. Upon closer investigation of the dyads that involved “squeezing” demonstrators only, demonstrators paired with observers that failed to learn tended to open fewer boxes, but this difference was not significant. There was also no difference between these dyads and those that included a “staggered-pushing” demonstrator (one-way ANOVA, F=2.446, p=0.129; Table 2; Fig. 2A).

**Figure 2.**
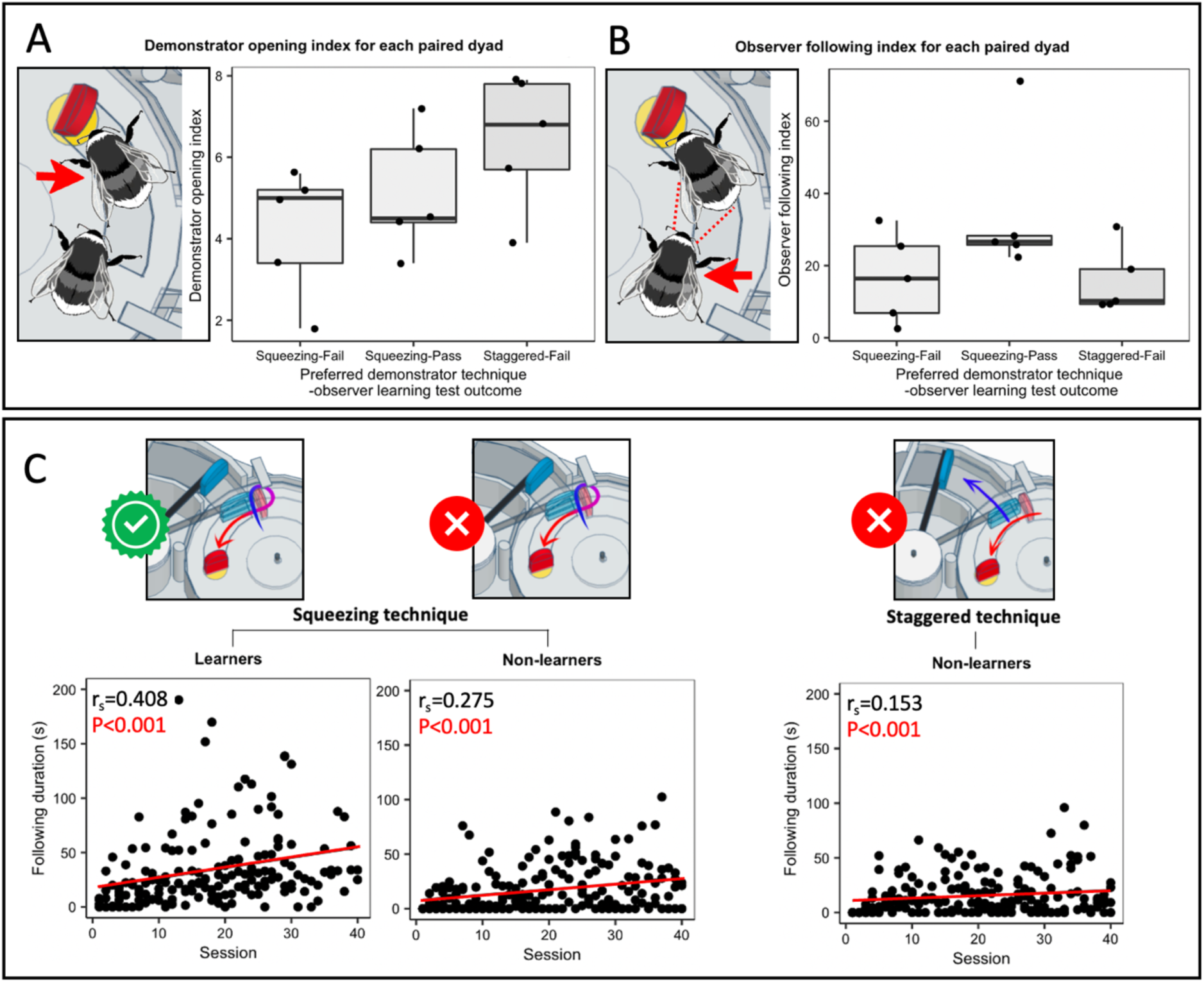
Demonstrator action patterns impacted the acquisition of two-step box-opening by observers. (A) Demonstrator opening index. The demonstrator opening index was calculated for each dyad as the total incidence of box-opening by the demonstrator / number of joint foraging sessions. (B) Observer following index. Following behaviour was defined as the observer being present on the surface of the box, within a bees’ length of the demonstrator, while the demonstrator performed box opening. The observer following index was calculated as the total duration of following behaviour / number of joint foraging sessions. Data in (A) and (B) were analysed using one-way ANOVA and are presented as boxplots, showing the median value and interquartile range. (C) Duration of following behaviour over the paired dyad joint foraging sessions. Following behaviour significantly increased with the number of joint foraging sessions, with the sharpest increase seen in dyads including a “squeezing” demonstrator and an observer that successfully acquired two-step box opening. Data were analysed using Spearman’s rank correlation coefficient tests. Individual data is presented in Supplementary Fig. 1.

**Table 2.**
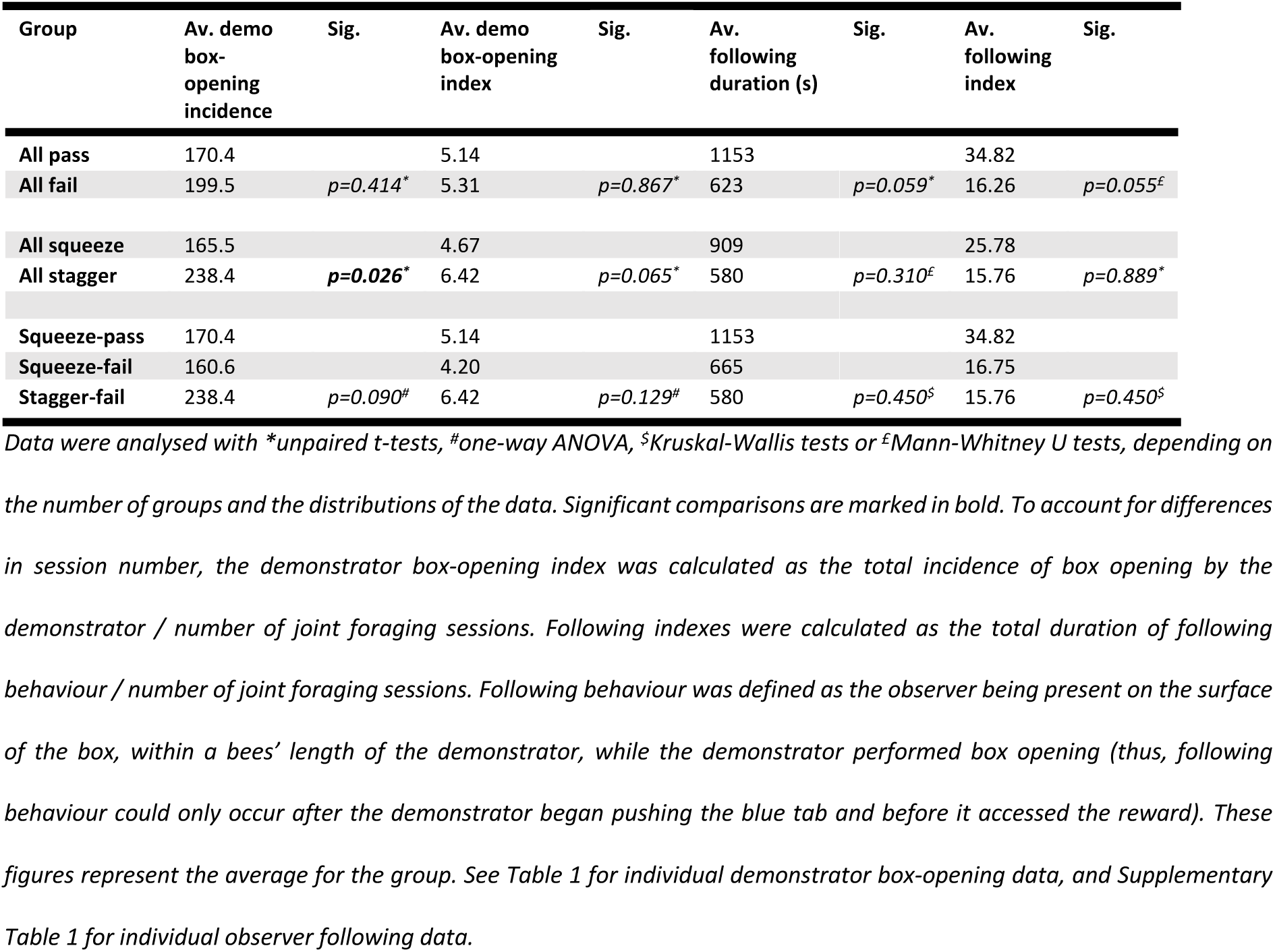
Paired dyad demonstrator and observer characteristics.

Furthermore, successful learners appeared to acquire the specific technique used by their demonstrator: in all cases, this was the “squeezing” technique. In the solo foraging sessions recorded for successful learners, these also tended to preferentially adopt the “squeezing” technique (Table 1). In short, being exposed to instances of two-step box-opening via “squeezing” appeared to be critical for successful transmission of the behaviour, and this variant was then repeated by learners.

### Observer following behaviour increased over time during the paired dyad experiments

To determine whether observer behaviour might have differed between those who passed and failed, we investigated the duration of their “following” behaviour, which was a distinctive behaviour that we identified during the joint foraging sessions. Here, an observer followed closely behind the demonstrator as it walked on the surface of the box, often close enough to make contact with the demonstrator’s body with its antennae. In the case of “squeezing” demonstrators, which often made several loops around the red tab, a following observer would make these loops also. To ensure we quantified only the most relevant behaviour, we defined following behaviour as “instances where an observer was present on the box surface, within a single bees’ length of the demonstrator, while it performed two-step box-opening”. Thus, following behaviour could be recorded only after the demonstrator began to push the blue tab, and before it accessed the reward. This was quantified for each joint foraging session for the paired dyad experiments (Supplementary Table 1). There was no significant correlation between the demonstrator opening index and the observer *following index* (Spearman’s rank correlation coefficient, rs=0.173, p=0.537; Supplementary Fig. 2), suggesting that increases in following behaviour were not simply due to there being more demonstrations of two-step box-opening available to the observer.

There was no statistically significant difference in the *following index* between dyads with “squeezing” and “staggered-pushing” demonstrators, between dyads where observers passed or failed, or when both demonstrator preference and learning outcome were accounted for (Table 2). This may have been due to the limited sample size. However, the *following index* tended to be higher in dyads where the observer successfully acquired two-step box-opening compared with those where the observer failed (34.82 vs. 16.26, respectively; Table 2) and in dyads with “squeezing” demonstrators compared with “staggered-pushing” demonstrators (25.78 vs. 15.76, respectively; Table 2). When both factors were accounted for, following behaviour was most frequent in dyads with a “squeezing” demonstrator and an observer that successfully acquired two-step box-opening (34.82 vs. 16.75 (“squeezing/fail” group) vs. 15.76 (“staggered-pushing/fail” group); Table 2).

There was, however, a strong and significant positive correlation between the duration of following behaviour and joint foraging session number, which equated to time spent foraging alongside the demonstrator. This association was present in dyads from all three groups but was strongest in the “squeezing/pass” group (Spearman’s rank order correlation coefficient, rs=0.408, p<0.001; Fig. 2C). This suggested that, in general, either the latency between demonstration commencement and observer following behaviour decreased over time, or that observers continued to follow for longer once arriving. However, the observers from the “squeezing-pass” group tended to follow for longer than any other, and the duration of their following increased more rapidly. This indicates that following a conspecific demonstrator as it performed two-step box-opening (and, specifically, via “squeezing”) was important to the acquisition of this behaviour by an observer.

### Bumblebees were unable to solve the two-step puzzle box without a demonstrator

In our previous study, several bees successfully learned to open the two-option, single-step box under “control” open diffusion conditions, which were conducted in the absence of a trained demonstrator across 6-12 days^18^. Thus, to determine whether the two-step box could be solved individually, we sought to conduct a similar experiment. Two colonies (C1 and C2) took part in these “control” open diffusion experiments. In brief, on 12 consecutive days, bees were exposed to open two-step puzzle boxes for 30 min pretraining and then closed boxes for 3 h. No trained demonstrator was added to the group. On each day, bees foraged willingly during the pretraining, but no boxes were opened in either colony during the experiment. While some bees were observed to probe around the components of the closed boxes with their proboscises, particularly in the early diffusion sessions, this behaviour generally decreased as the experiment progressed. A single blue tab was opened in full in colony C1, but this behaviour was neither expanded upon nor repeated. Considering that bumblebees primarily learn through association with either reward or punishment, and there was nothing to be directly associated with this action other than expended effort, this outcome is unsurprising.

### Observations from developing the demonstrator training protocol provided supportive evidence to suggest that bumblebees are unlikely to be able to acquire two-step box-opening individually

Learning to open the two-step box was not trivial for our demonstrators, with the finalised training protocol taking ∼2 days for them to complete (compared with several hours for our previous two-option, single-step box^18^). Developing a training protocol was also challenging. Bees readily learned to push the rewarded red tab, but not the unrewarded blue tab, which they would not manipulate at all. Instead, they would repeatedly push against the blocked red tab before giving up. This necessitated the addition of a temporary yellow target and reward beneath the blue tab, which in turn required the addition of the extended tail section (as seen in Fig. 1A), as during later stages of training this temporary target had to be removed and its absence concealed. This had to be done gradually and in combination with an increased reward on the final target, as bees quickly lost their motivation to open any more boxes otherwise. Frequently, reluctant bees had to be “coaxed” back to participation via provision with fully-opened boxes that they didn’t need to push at all. In short, bees seem generally unwilling to perform actions that are not directly linked to a reward, or that are no longer being rewarded. Notably, during the joint foraging sessions, demonstrators frequently pushed against the red tab before attempting to push the blue, even though they were able to perform the complete behaviour (and subsequently did so). The combination of having to move *away* from a visible reward and take a non-direct route, and the lack of any reward in exchange for this behaviour, suggests that two-step box-opening would be very difficult, if not impossible, for a naïve bumblebee to discover and learn for itself.

## Discussion

In this article, we present evidence suggesting that *Bombus terrestris*, a social invertebrate, is capable of learning a novel behaviour from a conspecific that cannot be learned through individual trial-and-error. Two-step box-opening involves an initial, unrewarded step, where bees must push a blue tab away from the path of a red tab, before pushing the red tab to receive a reward. This behaviour was so challenging that, unless an extra reward was added beneath the blue tab, demonstrators failed to learn two-step box-opening during training. This additional reward had to be removed gradually during later training stages to avoid bees refusing to open more boxes. Even so, 5/15 naïve observers successfully acquired the complete behaviour from the trained demonstrators. The fact that any observer bee was able to learn the complete two-step behaviour was striking precisely *because* they acquired the complete behaviour: these bees had never been exposed to any form of puzzle box, had not learned either of the two steps before being exposed to the other, and unlike the demonstrators had never been rewarded for pushing the blue tab. Yet, they were able to acquire the entire behaviour sequence via social learning.

In contrast, in a recent study, great tits were challenged with a two-step puzzle box task after they had previously learned one of the two steps^21^. While the birds could combine the two behaviours to solve the puzzle, they did not learn the complete two-step behaviour from the demonstrator: rather, they learned component behaviours and recombined them individually to form the full solution. The authors hypothesised that great tits might need both steps to be rewarded, at least initially, which mirrored the observations we made while developing our demonstrator training protocol. But bees in our study were still able to learn the complete behaviour without any reward for the first step or experience with any type of puzzle box, purely via exposure to a knowledgeable conspecific.

The results of the open diffusion control experiments, in which bees were exposed to puzzle boxes for 36 h across 12 days, were equally telling. No bee came close to opening even a single box and their interest in the closed boxes plummeted with time, although they continued to forage on opened boxes during pre-training. For comparison, in our previous work using the two-option puzzle box requiring only a single-step action, the two 12-day control colonies generated 13 learners between them in the absence of any demonstrator^18^. This remarkable capacity for innovation is consistent across paradigms: in previous work on string-pulling, some bumblebees were also reported to learn to pull strings without a demonstrator^15^. The failure of bees to open the two-step box independently, then, is unlikely to be due to a lack of behavioural flexibility. However, both of these previous examples involved single-step, directly-rewarded tasks. If bumblebee learning primarily relies on reward/punishment-based associations, it may simply not be possible for them to learn unrewarded actions, unless these are somehow linked to a rewarded action. This may be why bees succeeded in learning two-step box-opening only when exposed to a demonstrator performing the “squeezing” behavioural variant: this action pattern essentially combined the two steps into one, reducing the degree of temporal and spatial separation between the first step of the behaviour and the reward when compared with “staggered-pushing”. This may have permitted bees to form an association between the two. But the presence of the demonstrator itself was also key: observers closely “following” behind the demonstrator as it performed the key “squeezing” action essentially saw them “squeezing” too, accurately replicating the demonstrator’s behaviour without the need for intentional imitation. While there was no significant difference, the duration of “following” behaviour was also markedly increased among learners compared with non-learners, suggesting that this facilitated the transmission of two-step box-opening to an extent. Taken together, bees seem highly unlikely to be capable of solving the two-step puzzle box through individual learning, even though they were capable of learning to do so socially. While this capacity is impressive, it becomes more so when one considers that the ability to learn such behaviours is a key prerequisite for *cumulative culture*.

Cumulative culture refers to behavioural traditions that are acquired through social learning, and have (via the incorporation of novel innovations) become too complex or elaborate for an individual to re-innovate without any social input^1–3^. Often, this definition includes the stipulation that independent re-innovation must be impossible *within an individual’s own lifetime*. This has resulted in a dearth of laboratory-based experimental assessments of cumulative culture in animals. It is difficult to conceive of an experimental design that might convincingly demonstrate this capacity in long-lived species, such as primates, cetaceans or corvids^22^, but these are the species we tend to assume are most likely to be capable of this feat. Food washing behaviours by macaques^9^, *Pandanus* leaf tool designs by New Caledonian crows^12^ and the songs of humpback whales^23^ have all been proposed as potential examples of cumulative culture, but none have been confirmed through laboratory-based experiments. In the case of cetaceans, there are additional physical constraints: it is hard to see how one would even begin to approach such an experiment in a wild humpback whale. This does not mean that these animals are incapable of cumulative culture, or even that these examples do not represent it: it simply means that we cannot know for *sure* whether they do, and this allows room for a hard stance against it on the principle of parsimony. Opposing arguments also often fail to recognise that a single animal performing a behaviour once, potentially by chance during trial-and-error-learning, does not constitute solid evidence against cumulative culture. It does not mean that this behaviour has been properly incorporated into an individuals’ repertoire or that it will ever be repeated in a meaningful way again, which will of course stall any potential transmission to conspecifics via social learning. Certainly, it should not be taken as evidence that an otherwise population-wide, socially acquired behaviour is *not* the result of cumulative culture.

In this study, we did not aim to examine the maintenance of two-step box-opening as a behavioural tradition by a group of bumblebees over time, as we did in our previous work with the two-option box^18^. However, the evidence gathered by the present study and our previous work suggests that this would at least be cognitively plausible. This is especially notable as *Bombus terrestris* has not been confirmed to display any form of culture in the wild, cumulative or otherwise, although nectar-robbing and its handedness may represent strong candidates^9,30^. In fact, on the surface, this species seems vanishingly unlikely to show cumulative culture at all in the wild. Bumblebee colonies in temperate regions do not typically persist beyond a single biological generation^26^. They are survived by naïve queens who might have little opportunity to learn from experienced individuals, which would effectively reset any accumulation of complexity or efficiency. Still, in our study, bumblebees were able to learn a behaviour from a demonstrator that was complex enough that they failed to re-innovate it independently. This suggests that either the diminutive brains of bees are efficient beyond our current understanding, that this achievement is not so cognitively complex as previously assumed, or some combination of the two. Perhaps, then, the apparent lack of cumulative culture among wild bumblebees is as simple as a lack of opportunity and need^27^. Wild bees are perhaps unlikely to stumble across natural analogues of multi-step puzzle-boxes that they must solve in order to feed. But the question remains: *why* are they capable of such a feat, if it is not something necessary in their natural lives?

Social insects have some of the richest, most intricate behavioural repertoires in the entire animal kingdom. Their nest architectures are orders of magnitude larger than any individual, and are built in common. Leafcutter ants farm fungus on the leaves that they collect^28^, and honeybees communicate the distance and direction of resources via their dance language^29^. This behaviour was all once thought purely instinctive. However, we are increasingly beginning to appreciate the role of social learning in such behaviour: at least some components of the honeybee dance language appear to be shaped via social influences^30^. Some social insect species form colonies that last for years or even decades: these include honeybees^31^, tropical bumblebees^32 33 34^ and stingless bees^35,36^. If the learning abilities of these species resemble those of *Bombus terrestris*, these may be the best candidates where one might observe the natural occurrence of culture, and even cumulative culture. But even if behaviour relies on an innate, genetic basis in the present day, it is possible this was not always the case, as suggested by Waddington’s theory of genetic assimilation^37^ and Baldwin’s phenotype-first theory of evolution^38^. In particular, the latter theory suggests that beneficial learned behaviours can become increasingly instinctive as selection favours enhanced learning abilities or behavioural biases, reducing the time and energy needed to acquire the behaviour. Perhaps, then, it is even possible that the “ratchet” thought to characterise cumulative culture does not have to involve a high-fidelity learning mechanism such as imitation at all. If the ratchet must hold behaviours within a population for long enough and with sufficient fidelity for them to be improved upon at a later point, it is hard to imagine a maintenance system more accurate than genetic code. Perhaps the reason we have so often failed to see evidence of culture in non-human animals is that we have simply been looking too late. Bumblebees and their fellow social insects represent exciting models through which these questions can be pursued and, perhaps, also answered.

## Materials and methods

### Animal model

Bumblebees (*Bombus terrestris audax*) were obtained from Agralan, Ltd. (Swindon, UK). Whole colonies were transferred to 30.0 ξ 14.0 ξ 16.0 cm bipartite wooden nest boxes upon delivery, where they were housed for the duration of the experiment. Colonies were maintained at room temperature, and experiments were conducted under standardised artificial lights (12:12, high-frequency fluorescent lighting; TMS 24F lamps with HF-B 236 TLD [4.3 kHz] ballasts [Koninklijke Phillips NV, Amsterdam, Netherlands], fitted with Activa daylight fluorescent tubes [OSRAM Licht AG, Munich, Germany]). Bees foraged *ad libitum* on 20% w/w sucrose solution, which was provided via mass feeders overnight in the flight arena, with pollen provided directly to the nest box every two days. All experiments were conducted in accordance with the ASAB/ABS guidelines for the use of animals in research. There were no license or permit requirements for the experiments presented in this paper.

### Experimental set-up and puzzle box design

#### Puzzle box

The two-step puzzle box was a modified version of the two-option puzzle box used in previous work by our group^18^. This incorporated a transparent lid that rotated anti-clockwise around a central axis when a red tab was pushed, exposing a 50% w/w sucrose solution reward on a yellow target; however, an additional blue tab initially blocked the path of the red tab (Fig. 1A). Two stoppers prevented either tab from being moved in the incorrect direction, and a plastic “shield” around the box prevented bees reaching with their proboscis from the side of the box to obtain the reward without manipulating the tabs.

#### Flight arena

All experiments were conducted in a flight arena (66.0 × 60.0 × 30.0 cm) which was connected to the hive boxes with transparent acrylic tunnels (26.0 × 3.5 × 3.5 cm). Plastic strips used as sliding “doors” along this tunnel controlled access to the flight arena. Flaps cut into the arena sides allowed the removal and replacement of puzzle boxes with minimal disturbance to the bees, and brush strips lining the sides of the flight arena prevented bees from escaping during this process. The interior walls were lined with laminated paper displaying a random distribution of pink dots. The top of the flight arena was a sheet of transparent UV-transmitting Perspex acrylic, and cameras (iPhone 6S; Apple, Inc., Cupertino, CA, USA) were placed on top of the arena to record all experiments from above.

#### Individual tagging

Bees were marked with numbered Opalith tags (Opalithplättchen; Warnholz & Bienenvoigt, Ellerau, Germany) for individual identification^18^. Briefly, small groups of bees were allowed into the flight arena to forage on lidless puzzle boxes, with yellow targets (carrying 10 μl 50% w/w sucrose solution rewards) fully exposed and accessible. Bees that foraged from multiple boxes (i.e., were motivated foragers) were captured and tagged before being returned to the hive box. Bees were never used for experiments on the same day that they were tagged to prevent any confounding stress-associated effects, and to allow acclimation to the tag.

### Demonstrator training protocol

#### Manual stepwise training of two-step box opening

Potential demonstrators were identified during group foraging on lidless boxes, as described above. When a tagged bee was observed repeatedly and reliably coming back and forth between the nest box and the flight arena to forage, it was selected for further training, and all other bees were restricted to the nest box. Full details of the stepwise protocol can be found in the **Supplementary information**. Notably, to train bees to push the unrewarded blue tab as the first step of the behaviour, it was necessary to introduce a temporary additional reward beneath this tab, which was then removed during later training stages. Successful acquisition of two-step box opening was confirmed via an unrewarded learning test.

#### Demonstrator unrewarded learning test

Once a bee reliably opened two-step puzzle boxes in exchange for no reward beneath the blue tab and for 20 μl 50% w/w sucrose solution beneath the red tab, it proceeded to the unrewarded learning test. The full unrewarded learning test protocol can be found in the **Supplementary information**. The average training period for a wholly naïve bee was two days and, in an attempt to reduce this time, three trained demonstrators (A-17, A-24 and C26) were originally used as observers in the paired dyad experiments before being put through the stepwise training protocol and the unrewarded learning test.

A total of 13 bees from four colonies (colony IDs A-D) passed the test and were used as demonstrators in the paired dyad experiments. Three of these (A-8, A-27 and A-6) were used to pilot these experiments, and one (B-25) died before it could be used as a demonstrator. Thus, nine bees in total were used as demonstrators for the paired dyad experiments (Table 1).

### Paired dyad experiments

*Observer selection*. In total, 15 observers from three colonies took part in these experiments. Each observer was paired with a trained demonstrator, with some demonstrators being used for multiple observers in succession (Table 1). Observers were selected in the same manner as demonstrators during group foraging on lidless boxes, with only the most motivated and reliable foragers chosen. Thus, while observers were familiar with the yellow target indicating the presence of a reward, they had no experience with closed boxes, or with the movement of the tabs. They also never had any experience with being rewarded for pushing the blue tab, as demonstrators did during early training phases.

#### Joint foraging phase

Prior to each joint foraging session, the demonstrator and the observer were held in the tunnel for 3 min, starting from when both were present. Following this, they were released into the flight arena and presented with three closed puzzle boxes, each filled with 20 μl 50% w/w sucrose solution. As these were opened, they were removed from the arena, cleaned with 70% ethanol to remove olfactory cues, refilled and replaced. Once the demonstrator returned to the tunnel (or after 20 min elapsed, at which point the demonstrator was manually returned to the nest box) the experimenter removed one box, opened it, and placed it back in the arena so the observer could access the reward. This was again intended to preserve foraging motivation. Occasionally, demonstrators lost motivation to forage during this phase: to counter this, if they ever failed to open more than two boxes in a single session, they were given individual foraging sessions until they opened boxes consistently again.

#### Observer unrewarded learning test

After 30 joint foraging sessions, the observer proceeded to an unrewarded learning test in the absence of the demonstrator. This was identical to the test used for demonstrators, but if the box was opened in the 15 min time limit, the observer was given a yellow acrylic chip loaded with 10 μl 50% w/w sucrose solution followed by a closed box with *50* μl 50% w/w sucrose solution on the target. The observer was then allowed to forage *ad libitum* on closed puzzle boxes (containing the usual 20 μl 50% w/w sucrose solution), until it either returned to the nest box or stopped opening boxes for 3 min. Observers who passed proceeded to the solo foraging phase, while those that failed returned to joint foraging sessions with the demonstrator. Additional learning tests were given after every five “remedial” joint foraging sessions, but if the observer failed three times it was considered to have failed to acquire two-step box-opening. At this point, joint foraging was ceased.

#### Solo foraging phase

Observers that passed the learning test proceeded to the solo phase. Here, it was challenged with 10 foraging sessions in the absence of the demonstrator. Each session was preceded by a 3 min waiting period in the tunnel and then lasted up to 10 min, following which the bee was manually returned to the nest box if it had not already done so itself. Three puzzle boxes, each containing 50 μl 50% w/w sucrose solution, were placed in the arena, and if none were opened during a session, the bee was given a yellow acrylic chip with 10 μl 50% w/w sucrose solution before being returned to the nest box. This was done in an effort to maintain their foraging motivation for further sessions.

### Control open diffusion experiments

Two colonies (IDs C1 and C2) were used for these experiments, which followed a similar protocol as the control open diffusion experiments described in our previous work^18^. These experiments aimed to determine whether bees could spontaneously learn to open the two-step box during an extended period of exposure. Multiple bees managed to open a single-step puzzle box without a demonstrator in our previous experiments^18^.

In brief, each day at ∼9.30 am, the mass feeders were removed from the flight arena and bees were returned to the nest box. If more than two honeypots were full, the sucrose solution was removed using a handheld pipette, with care being taken to avoid damaging any part of the hive structure. This ensured a strong motivation to forage. After ∼30 min, bees were allowed unrestricted access to the flight arena, where they received 30 min group pre-training with eight lidless boxes, with the yellow targets (bearing 10 μl w/w 50% sucrose rewards) fully exposed. This daily pre-training ensured that the bees acquired maintained a strong association between the colour yellow and the reward, and encouraged as many bees into the flight arena as possible before the experiment began. Following the pre-training, the boxes were removed, wiped with 70% ethanol, refilled with 20 μl w/w 50% sucrose, closed, and replaced to begin the diffusion experiment. Each diffusion session lasted for 3 h.

### Video analysis

All videos were analysed using BORIS 7.10.2, which permitted the coding of point events, the extraction of event durations, and the assigning of each event to a bee ID, box ID, tab colour and opening method as appropriate^39^.

*Paired dyad experiments*. For both the joint sessions and post-learning solo foraging sessions, point events were coded when boxes were opened (“demonstrator opening incidence” in the joint sessions, “learner opening incidence” in the solo sessions). The start and end of observer “following behaviour” was also coded as point events in the joint sessions. Following behaviour was defined as the observer being present on the surface of the puzzle box, within a bees’ length of the demonstrator, while it performed box-opening (i.e., following behaviour could only occur after the demonstrator began pushing the blue tab and before it accessed the reward). The total duration of following behaviour in each joint session was thus extracted.

As the dyads underwent different numbers of joint sessions, depending on whether the observer acquired two-step box-opening or not, demonstrator opening incidence and observer following duration were indexed to permit statistical comparisons. The demonstrator opening index for each dyad was calculated as the total incidence of box-opening / the number of joint sessions, and the following index were calculated as the total duration of following / the number of joint sessions.

#### Box opening method

As aforementioned, two different methods to open the two-step puzzle box were identified during these experiments: “squeezing” and “staggered-pushing” (Fig. 1B). Thus, all incidences of box-opening in both the paired dyad and open diffusion experiments were additionally labelled with the box-opening technique that was used. As in some cases, bees would leave the box surface during the opening or move between the blue and red tabs several times, even while performing the key squeezing movement between the tabs (sometimes repeatedly), any openings where this squeezing action was incorporated were labelled “squeezing”.

### Statistical analysis

Data were analysed using R 4.0.4^40^. Comparisons between two groups were made with unpaired t-tests or Mann Whitney U tests, depending on the normality of the data. Comparisons between more than two groups were conducted with one-way ANOVA or Kruskal-Wallis tests, as appropriate. Correlation analysis was performed using Spearman’s rank order correlation coefficient tests. p<0.05 was considered to indicate a statistically significant difference.

## Author contributions

**Alice D. Bridges:** Conceptualization, Methodology, Supervision, Investigation (video analysis), Formal Analysis, Visualization, Writing (original draft preparation) **Amanda Royka:** Methodology, Investigation (paired dyad experiments, control diffusion experiments, video analysis) **Tara Wilson:** Investigation (paired dyad experiments, control diffusion experiments, video analysis) **Lars Chittka:** Funding Acquisition, Conceptualization, Supervision, Resources, Writing (review & editing)

## Additional information

Supplementary information is available for this paper.

## Supplementary information

### Supplementary figures

**Supplementary Figure 1.**
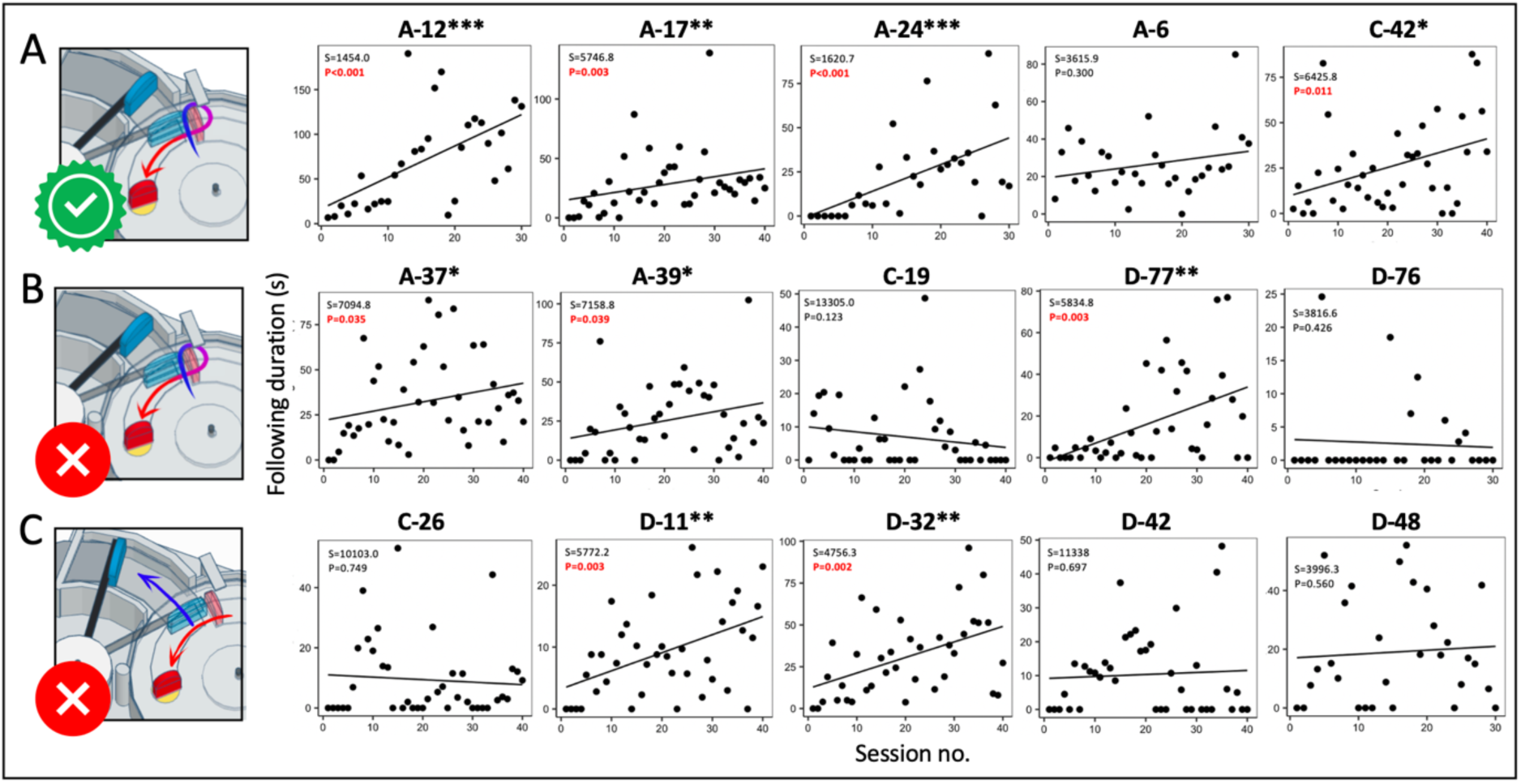
Following duration over the paired dyad joint foraging sessions for individual paired dyads. (A) Dyads where the observer acquired two-step box-opening and the demonstrator preferred the squeezing technique. **(B)** Dyads where the observer failed to acquire two-step box-opening and the demonstrator preferred the squeezing technique**. (C)** Dyads where the observer failed to acquire two-step box-opening and the demonstrator preferred the squeezing technique. Data were analysed using Spearman’s rank correlation coefficient tests, and significant results are highlighted in red. Graph titles refer to the observer ID.

**Supplementary Figure 2.**
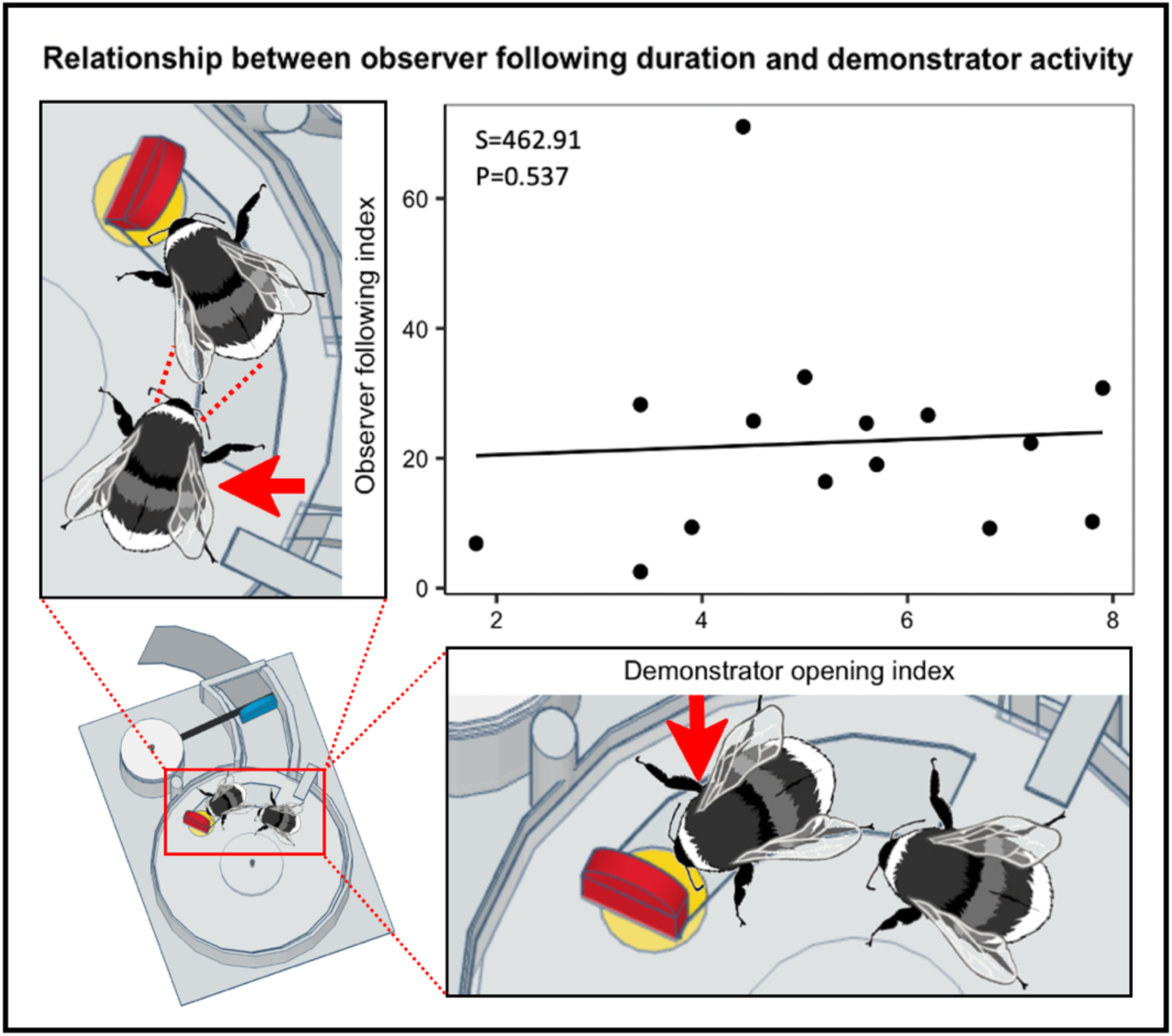
There was no significant correlation between demonstrator opening index and observer following index in the paired dyads. This suggested that increases in following behaviour were not simply due to there being more demonstrations of two-step box-opening available to the observer. To account for differences in session number, demonstrator box-opening indexes were calculated as the total incidence / number of sessions. Following indexes were calculated as the total duration of following behaviour / number of sessions. Data were analysed using a Spearman’s rank correlation coefficient test.

**Supplementary Figure 3.**
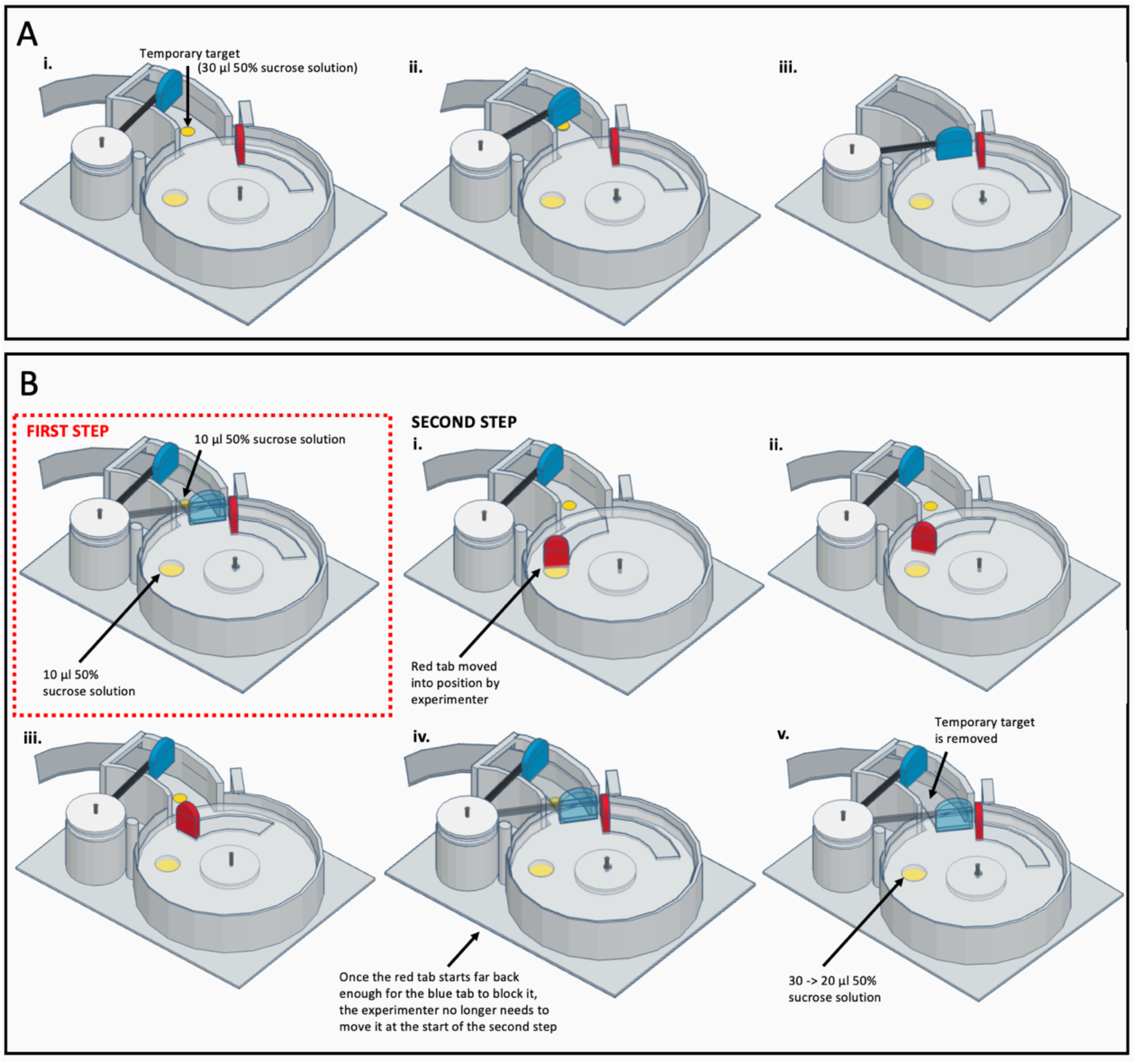
Stepwise demonstrator training. (A) Training a demonstrator to push the blue tab. A temporary yellow target bearing 30 μl 50% sucrose solution was added to the tail section of the box; the blue tab was initially positioned so that this was fully exposed (i). Once the bee learned the location of the reward, the blue tab was moved further over the target. Initially, the reward could still be obtained by reaching under the tab, but the tab was often pushed forward as the bee attempted this (ii). This continued until the reward was inaccessible without pushing the tab, and the blue tab blocked the red tab and prevented it from being moved (iii). **(B)** Training a demonstrator to push the red tab. Yellow targets now bore 10 μl 50% sucrose solution. When the bee was reliably pushing the blue tab from the fully-closed configuration (A iii), phase 2 of training began. Bees continued to push the blue tab and obtain the reward beneath (first step; see red inset panel); once completed, the experimenter used tweezers to move the red tab forwards and expose the second yellow target with reward (second step; i). As above, the location of the red tab in the second step was progressively shifted so that the second yellow target was increasingly inaccessible (ii-iii). Once the bee was able to push the red tab from a position far enough back that the blue tab blocked it when closed, the experimenter no longer intervened before the step 2 (iv). The final step was the removal of the temporary yellow tab beneath the blue door and a temporary increase of the reward to 30 μl 50% sucrose solution, decreased to 20 μl once the bee reliably opened the box by pushing both tabs with no reward beneath the blue tab. Once it reliably did this for 20 μl, it progressed to the learning test.

### Supplementary tables

**Supplementary Table 1.**
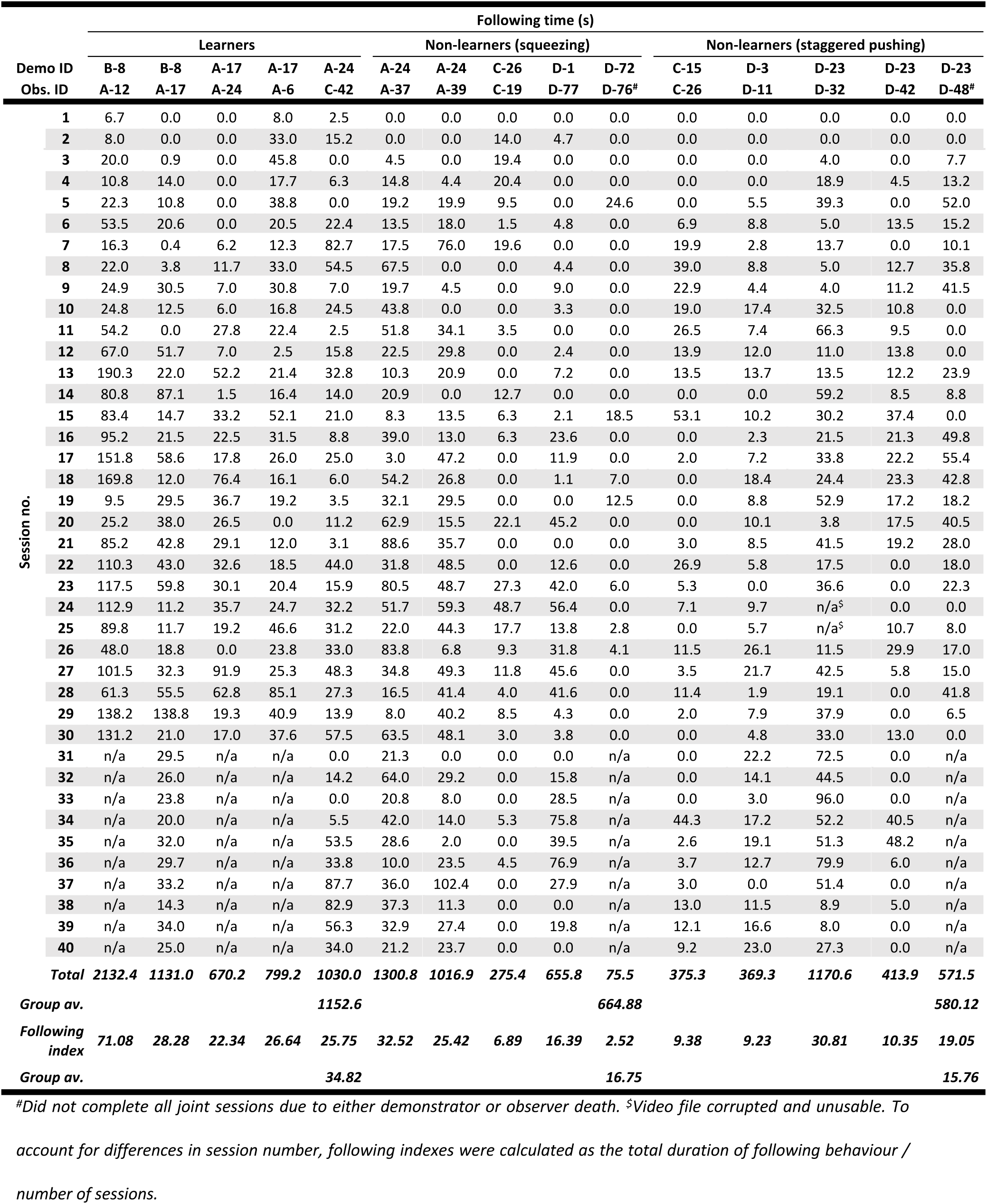
Duration of observer following behaviour in the paired dyad experiments.

### Supplementary methods

#### Puzzle box design elements

Box bases were 3D-printed to ensure consistency, and the surface was covered in laminated grey RGB-neutral paper (hex #555555), which was lightly sanded to provide grip. A Petri dish lid served as the rotating lid of the box, with one cut-out track available for bees to use. A second cut-out track was made in the lid for balance but was covered with a thin sheet of clear plastic. White acrylic “washers” and nails formed the two rotating mechanisms. The red and blue tabs were cut from 2 mm-thick polyethylene craft foam sheets, and while the red tab was affixed to the front of the cut-out track on the lid, the blue tab was attached to a thin strip of plastic. The other end of this plastic “arm” was incorporated into the rotating mechanism on the column. An opaque cover, made from laminated grey paper, was cut so that it would follow the curve of the tail section and was affixed to the bottom of the blue tab, as depicted. The yellow target indicated the location of the reward (50% w/w sucrose solution) and was always visible, but inaccessible without opening the box. The outer “shield” surrounding the box prevented bees obtaining rewards by squeezing under the lid from the sides, and two stoppers prevented the movement of either tab in inappropriate directions.

#### Stepwise demonstrator training protocol

The stepwise demonstrator training protocol is presented in Supplementary Figure 3. During training, all boxes were washed with 70% ethanol after depletion and before replacement, to ensure the removal of any olfactory cues. The first stage of training (Supplementary Fig. 3A) involved training the demonstrator to push the blue tab out of the way of the red tab: during this stage, a temporary yellow target and sucrose solution reward was added beneath the blue tab.

Once a demonstrator could reliably access the initial reward beneath the blue tab (from Supplementary Fig. 3A, position iii) they proceeded to the second stage (Supplementary Fig. 3B). From this point, only one box was presented at a time to discourage bees from pushing multiple blue tabs without trying to push any red tabs. The initial reward was reduced from 20 to 10 μl 50% w/w sucrose solution, and an additional 10 μl 50% w/w sucrose solution reward was added to the final yellow target. Once the bee had obtained the reward beneath the blue tab, pushing it out of the way of the red tab in the process (Supplementary Fig. 3B inset), the experimenter used tweezers to manipulate the position of the red tab with respect to the second reward. As with the first reward, this was made progressively harder to obtain (Supplementary Fig. 3B i-iii), until the red tab was set far enough back that it was blocked by the blue tab and no longer required experimenter interference (Supplementary Fig. 3B iv).

At this point, the temporary reward beneath the blue tab was removed, but this had to be done slowly and progressively. When bees found no reward beneath the blue tab, they would soon refuse to open any more boxes, necessitating provision with boxes with the temporary target and reward reinstated (and easily accessible). Initially, one reward was removed per three boxes presented, then two per four boxes, then two per three boxes, until finally the temporary reward was removed entirely. The sucrose reward beneath the red tab also had to be tripled to 30 μl during this stage, or bees became reluctant to open more boxes. The final stage of training involved the reduction of this reward back to 20 μl, to ensure that demonstrators would not be satiated too quickly in subsequent experiments, and maximise the number of their demonstrations. If a bee stopped opening boxes at any point, it was presented with “easier” configurations from earlier training stages until it resumed its box-opening activity. Once a bee opened two boxes in the final configuration in exchange for 20 μl 50% sucrose solution, they proceeded to the unrewarded learning test.

#### Unrewarded learning test

Here, the bee was presented with a single, closed box with distilled water in place of the sucrose reward, to avoid any confounding olfactory cues. If the box remained unopened after 10 min had elapsed, a yellow acrylic chip bearing 10 μl 50% w/w sucrose solution was placed in the arena to retain foraging motivation. This chip was removed when depleted, and an additional 5 min were given to open the box. The increased duration of the test compared with that used for the two-option puzzle box task in our previous work^18^ was intended to reflect the increased difficulty of the two-step box. Bees that failed to open the box within the time limit returned to training until they met the unrewarded test criterion again. Bees that opened the box within the time limit ‘passed’ the test, and to prevent refusal to open further boxes in response to the distilled water, were immediately provided with a yellow acrylic chip carrying 10 μl 50% w/w sucrose solution. The experimenter replaced the opened box with a closed one containing 20 μl 50% w/w sucrose solution on the target, and the bee was then allowed to solve rewarded puzzle boxes *ad libitum* until it attempted to leave the flight arena.

